# Molecular characterization of carbapenem resistant *Klebsiella pneumoniae* clinical isolates: Preliminary experience from a tertiary care teaching hospital in the Himalayas

**DOI:** 10.1101/2021.09.10.459727

**Authors:** Mohit Bhatia, Varun Shamanna, Geetha Nagaraj, Dharmavaram Sravani, Pratima Gupta, Balram Ji Omar, Deepika Chakraborty, K. L. Ravikumar

## Abstract

**Introduction:** *Klebsiella pneumoniae* is recognized as an urgent threat to human health because of the emergence of multidrug-resistant (MDR) and hypervirulent strains. Development of novel highly effective and safe treatment options is the need of the hour. One of the ways of achieving this goal is by conducting molecular characterization studies of antibiotic resistant bacteria.

**Hypothesis/Gap Statement:** The government of India is committed to generate validated AMR data from different parts of the country. Efforts are being made to perform molecular characterization of MDR and potentially virulent bacteria by whole-genome sequencing (WGS). However, the data that we have at present is skewed as many parts of the country remain underrepresented. Uttarakhand is one such state located in the Himalayan belt of India, with relatively poor access to healthcare and a paucity of research.

**Aim:** This study was performed to generate WGS based preliminary data about the population structure, multi-locus sequence types (MLST), and virulence factors of CRKp isolates recovered from patients in a tertiary care teaching hospital and institute of national importance, located in Rishikesh, Uttarakhand, India.

**Methodology:** A cross-sectional study was conducted at a tertiary care teaching hospital. It included twenty-nine randomly selected and archived carbapenem resistant *Klebsiella pneumoniae* (CR-Kp) isolates obtained from various clinical samples submitted in the Bacteriology laboratory for culture and sensitivity testing, from July 2018 to August 2019. After preliminary identification (ID) and antibiotic susceptibility testing (AST), as per standard guidelines, these isolates were sent to Central Research Laboratory (CRL), Bengaluru, India, for further characterization & WGS.

**Results:** Twenty-seven out of twenty-nine test isolates were CRKp. Among the 27 CRKp isolates, ST14 was the most common sequence type (8, 29.6%), followed by ST231 (5, 18.5%) and ST147(3, 11.1%) respectively. KL2 (9/27, 33.3%) and KL51 (8/27, 29.6%) were dominant K loci types in this study. Out of 5 O antigens identified, O1 and O2 together accounted for 88.9% (n=27) CRKp isolates. Yersiniabactin and Aerobactin were identified in 88.9% (24/27) & 29.6% (8/27) of the CRKp isolates of the isolate. Regulatory genes rmpA2 and rmpADC were found in 14.8% (4/27) and 3.7% (1/27) isolates respectively. The predominant plasmid replicons present were ColKP3 (55.5%), IncFII(K) (51.8%), IncFIB(pQil) (44.4%), IncFIB(K) (37%), IncR (33.3%) and Col44 0I (18.5%) respectively. A perfect agreement (100%) was observed between phenotypic and genotypic resistance profiles in the case of fluoroquinolones, penicillins, Beta Lactam/Beta Lactam Inhibitor combinations (BL/BLI), and cephalosporins respectively. As compared to phenotypic resistance, higher genotypic resistance for aminoglycosides (96.3%) and folate pathway inhibitors (92.6%) respectively, was observed.

**Conclusion:** This study emphasizes the need for continued genomic surveillance of emerging CRKp and other MDR bacteria in Uttarakhand and neighbouring states of India. This in turn would help in generating critical information that can be used to assess the emergence, dissemination, and potential impact of important variants.

## Introduction

*Klebsiella pneumoniae* is recognized as an urgent threat to human health because of the emergence of multidrug-resistant (MDR) and hypervirulent strains associated with hospital outbreaks and severe community-acquired infections [1]. Of particular importance is carbapenem resistant *Klebsiella pneumoniae* (CRKp), which apart from being multi-drug resistant, is associated with high morbidity and mortality particularly among persons with prolonged hospitalization exposed to invasive devices [2].

Owing to paucity of treatment options for life threatening infections caused by MDR Gram negative bacteria (including CRKp), there is renewed interest in using highly toxic and narrow spectrum antibiotics like polymyxins, which were discovered almost six decades ago [3]. Development of novel highly effective and safe treatment options is the need of the hour. One of the ways of achieving this goal is by conducting molecular characterization studies of antibiotic resistant bacteria. In a review published by Karakonstantis et al, it has been highlighted that determination of genetic resistance mechanisms, particularly in CRKp isolates, may play a vital role in deciding novel treatment options like ceftazidime/avibactam, imipenem/relebactam etc., in hospitalised patients [4].

India has been referred to as the “Antimicrobial resistance (AMR) capital” of the world’ [5]. As per the ‘scoping report on antimicrobial resistance in India (2017)’, under the aegis of the Department of Biotechnology, Government of India, >70% isolates of *Klebsiella pneumoniae* were resistant to broad-spectrum antibiotics like fluoroquinolones and third generation cephalosporins. Increasing rates of resistance to carbapenem antibiotics and beta-lactam/beta-lactamase inhibitor combination antibiotics like piperacillin-tazobactam was also observed in *K. pneumoniae*, by the investigators [6]. The government of India is committed to generate validated AMR data from different parts of the country. Efforts are being made to perform molecular characterization of MDR and potentially virulent bacteria by whole-genome sequencing (WGS). These initiatives will significantly improve our ability to analyse outbreak scenarios and characterize the transmission dynamics & mechanisms of antibiotic resistance, of important pathogens [7]. However, the data that we have at present is skewed as many parts of the country remain underrepresented. Uttarakhand is one such state located in the Himalayan belt of India, with relatively poor access to healthcare and a paucity of research.

This study was performed to generate WGS based preliminary data about the population structure, multi-locus sequence types (MLST), and virulence factors of CRKp isolates recovered from patients in a tertiary care teaching hospital and institute of national importance, located in Rishikesh, Uttarakhand, India. This institute was established almost a decade ago and consistent efforts were being made to streamline the hospital infection control and antibiotic stewardship related activities, since it’s time of conception. A review of our partially documented cumulative antibiogram from January 2018 to December 2020 revealed, that 14.3% (279/1947) of MDR Gram negative bacterial isolates, were carbapenem resistant *Klebsiella pneumoniae*. Being classified as “Priority 1 pathogens” by the World Health Organisation (WHO) [8], CRKp isolates were selected to kick-start a genomic surveillance programme of MDR bacteria, at our institute. To the best of our knowledge, this study will be the first of its kind from the northern hilly terrain of India.

## Methods

### Bacterial isolates and phenotypic characterization

A cross-sectional study was conducted at a tertiary care teaching hospital, after obtaining ethical approval (AIIMS/IEC/18/477 & KIMS IEC/S12-2017). The study included twenty-nine randomly selected and archived carbapenem resistant *Klebsiella pneumoniae* (CR-Kp) isolates obtained from various clinical samples submitted in the Bacteriology laboratory for culture and sensitivity testing, from July 2018 to August 2019. After preliminary identification (ID) and antibiotic susceptibility testing (AST), as per standard guidelines [9,19], these isolates were sent to Central Research Laboratory (CRL), Bengaluru, India, for further characterization & WGS. At CRL, the test isolates were subjected to ID & AST, using the VITEK-2 compact system (bioMérieux), the results of which were interpreted as per Clinical Laboratory Standards Institute (CLSI) guidelines 2021 [11].

### Molecular characterization

Genomic DNA was extracted from bacterial isolates with Qiagen QIAamp DNA Mini kit, in accordance with the manufacturer’s instructions. Double-stranded DNA libraries with 450 bp insert size were prepared and sequenced on the Illumina platform with 150 bp paired-end chemistry. The genomes passing the QC were assembled using Spades v3.14 to generate contigs and annotated with Prokka v1.5 [12]. SNPs were identified for all 29 *K. pneumoniae* isolates by mapping of reads to the NCBI reference genome, *K. pneumoniae* strain NTUH-K2044, NC_006625.1 (https://www.ncbi.nlm.nih.gov/nuccore/NC_006625.1/). MLST, K-and O-antigen types, virulence factors and AMR genes were identified using the protocols as detailed in www.protocols.io [13].

## Results

1. Baseline information of the collection: The test isolates were obtained from clinical samples of patients with an age range of 16 months to 79 years (mean age of 42.81 years; male: female=23: 6), twenty-seven of which were CRKp. Most isolates were from urine (12/29; 41.4%) and pus (11/29; 37.9%). Sequencing results revealed that all test isolates were *K. pneumoniae* sensu stricto.
2. Clonal distribution: Among the 27 CRKp isolates, ST14 was the most common sequence type (8, 29.6%), followed by ST231 (5, 18.5%) and ST147(3, 11.1%) respectively. Sequence types 11, 15, 43, 395, 437 and 3392 together constituted 37% of the CRKp isolates. One novel ST (ST16-1LV) with XDR was identified in a CRKp isolate.

KL2 (9/27, 33.3%) and KL51 (8/27, 29.6%) were dominant K loci types in this study. The phylogenetic tree of the test isolates shows the correlation between capsular locus type and sequence type, for example of KL2 mainly with ST14, and of KL51 with ST231 and ST147 respectively. (Figure 1).

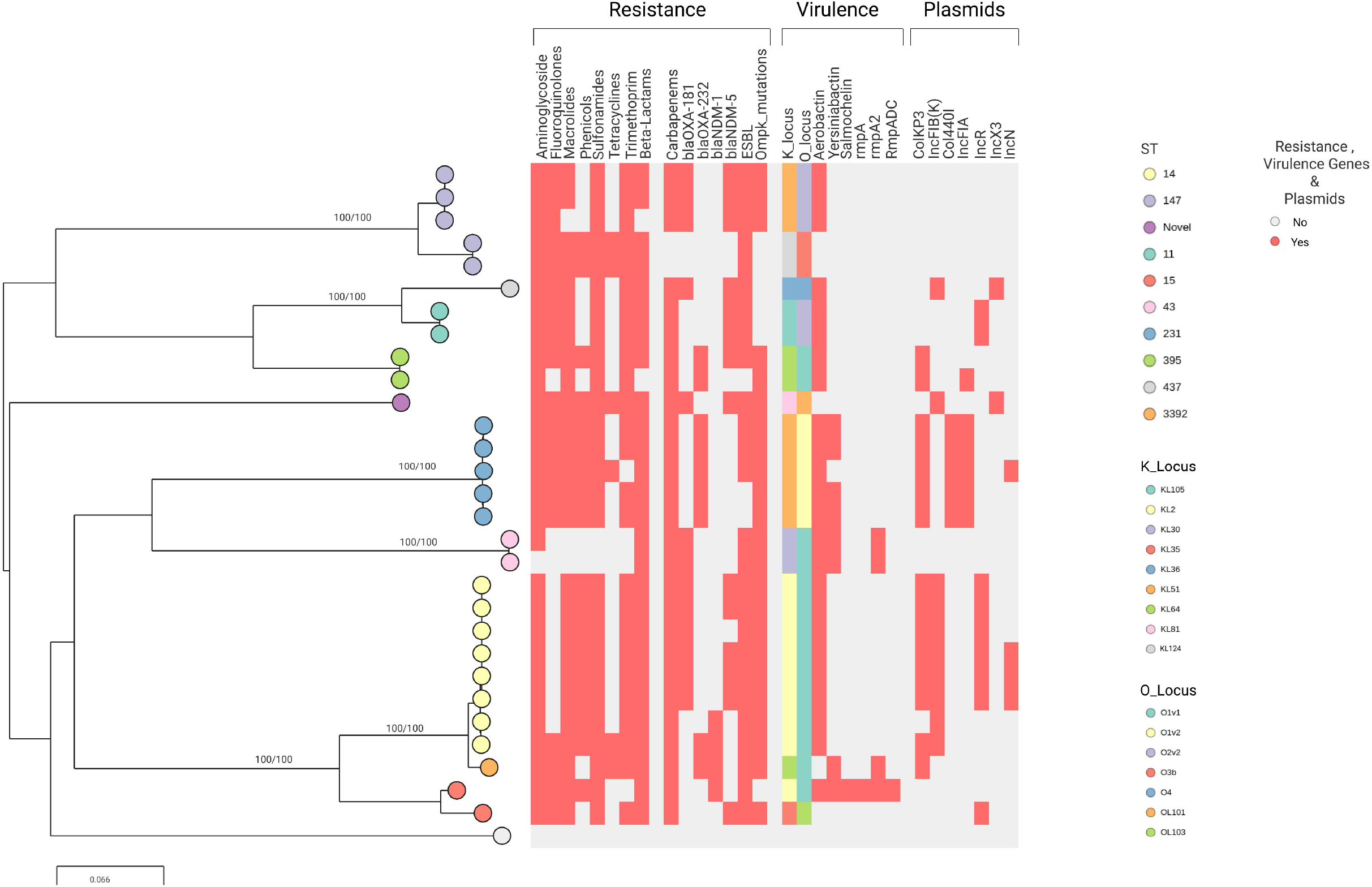

Out of 5 O antigens identified, O1 and O2 together accounted for 88.9% (n=27) CRKp isolates (Table 1). Other antigens present included O4 (3.7%), OL101 (3.7%), OL103 (3.7%). O-locus (OL) types also showed ST-specific distribution. In particular, O1v1 was found in STs 14, 43, 395, 15 and 3392, O1v2 in ST231 and O2v2 in STs 147 and 11 respectively (Table 1 & Figure 1). The two carbapenem sensitive MDR *K. pneumoniae* isolates belonged to ST 147, with K & O loci being KL124 and O3b, respectively.

3. Virulence factors & plasmid repertoire: Predominant virulence factor genes and plasmid replicons found in CRKp isolates in the present study, have been summarized in Table 1. Yersiniabactin, an iron uptake locus (ybt), was identified in 88.9% (24/27) of the isolates. Aerobactin, a critical virulence factor for Hypervirulent *K. pneumoniae* (HvKP), was found in 29.6% (8/27) of the isolates, all belonging to ST231, 43, 15 & 3392 respectively. Regulatory genes rmpA2 and rmpADC were found in 14.8% (4/27) and 3.7% (1/27) isolates respectively. While Colibactin and rmpA genes were absent in all isolates, Salmochelin was found in only one isolate.

The predominant plasmid replicons present in the 27 CRKp isolates were ColKP3 (55.5%), IncFII(K) (51.8%), IncFIB(pQil) (44.4%), IncFIB(K) (37%), IncR (33.3%) and Col44 0I (18.5%) respectively.

4. Antimicrobial resistance profile and their distribution: Table 2 depicts phenotypic and genotypic antimicrobial resistance profile of 27 CRKp test isolates. High phenotypic resistance rates to aminoglycosides (88.8%), fluoroquinolones (100%), penicillins (100%), BL/BLI (100%), cephalosporins (100%), folate pathway inhibitors (81.5%), and glycylcyclines (66.7%) respectively, were observed. Resistance to polymyxins was observed in 3.7% of the isolates. A perfect agreement (100%) was observed between phenotypic and genotypic resistance profiles in the case of fluoroquinolones, penicillins, Beta Lactam/Beta Lactam Inhibitor combinations (BL/BLI), and cephalosporins respectively. As compared to phenotypic resistance, higher genotypic resistance for aminoglycosides (96.3%) and folate pathway inhibitors (92.6%) respectively, was observed. While no colistin resistance genes were observed in 1 phenotypically resistant isolate, another isolate showed a disruption of the MgrB gene with a colistin MIC value in the intermediate range (≤0.5 µg/ml). None of the CRKp isolates contained tigecycline resistance genes.

All 27 CRKp isolates showed phenotypic resistance to imipenem, meropenem and ertapenem. Of these, 77.8% (21/27) carried *bla*_*OXA-48*_ like genes (*bla*_*OXA181*_ and *bla*_*OXA232*_), and 55.6% (15/27) carried *bla*_*NDM-1/5*_, respectively (Table 3). We observed that 71.4% (n=15/21) of the *bla*_*OXA-48*_ like genes carrying CRKp isolates harboured a plasmid with the ColKP3 replicon sequence (Figure 1). Notably, no *bla*KPC genes were detected.

In the present dataset, resistance to carbapenems was also conferred by ESBLs when combined with porin loss-of-function mutations. Disruption of the Ompk35 porin (n=21) and insertion mutations of TD/GD amino acids at position 115 of Ompk36 gene (n=19) were found in CRKp isolates. The disruption of major outer membrane protein genes (Ompk35/Ompk36) along with ESBLs (*bla*_*CTX-M-15*_, *bla*_*SHV-11*_, *bla*_*SHV-31*_, *bla*_*OXA-1*_, *bla*_*OXA-9*_, *bla*_*TEM-1*_, *bla*_*FONA-5*_, *bla*_*CMY-6*_, *bla*_*SFO-1*_) were detected in 40.7% of the isolates (11/27). There were 15 isolates in which multiple ESBL genes were detected.

## Discussion

In the present study, all clinical isolates were correctly identified as *Klebsiella pneumoniae* by VITEK-2 Compact, taking whole genome sequencing results as gold standard. Keeping in mind the potential clinical and epidemiological significance of each phylogroup, accurate identification of microbial isolates is imperative. Inability to distinguish species within the *K. pneumoniae* complex using phenotypic methods has been reported in the past by Bowers et al [14].

The predominant sequence types observed in our study were ST14, ST231 and ST147. A noteworthy finding was the absence of ST258, the most prevalent and extensively disseminated KPC-producing *K. pneumoniae* globally [15], in our collection. These findings are in concordance with published data from India [15]. ST14 and ST147 have often been described as international high-risk clones associated with extensive drug resistance (XDR) [16,17]. ST231 is an emerging CRKP epidemic clone which was first reported in 2013 in Delhi, India. This clone has also been reported from several other southeast Asian and European countries and has frequently been found to be MDR, combining resistance to carbapenems, extended-spectrum cephalosporins, and broad-spectrum aminoglycosides [15,18,19]. Another interesting finding of our study was that ST11 was identified in 7.4% of the CRKp test isolates. The ST11 clone is a single locus variant of ST258 and is associated with the spread of KPC in Asian countries like China and Taiwan [20]. Reports of presence of these clones in the Indian subcontinent [21], has made the AMR situation in the country, even more gruesome.

Owing to paucity of treatment options for infections caused by MDR *K. pneumoniae* strains, newer avenues in the form of lipopolysaccharide and capsule polysaccharide-based vaccines, are currently being explored [22]. Therefore, it is important to address the high diversity of surface-exposed polysaccharides, synthesized as O-antigens (lipopolysaccharide) and K antigens (capsule polysaccharide), present in *K. pneumoniae*. KL2, KL51 and O1, O2 were the predominant K-loci and O serotypes respectively, among CRKp isolates in our study. These findings are in concordance with a study conducted by Follador et al, in which serotypes O1, O2 and O3 were most prevalent in the sample set, accounting for approximately 80 % of all infections. In contrast, K serotypes showed an order of magnitude higher diversity and differed among infection types [22].

In the present study, virulence factor genes like yersiniabactin, aerobactin and carbapenemase carrying ColKP3 plasmid replicons were predominantly observed in ST14 & ST231 CRKp clones. Carriage of resistance markers on plasmids and virulence genes on integrative conjugative elements, is often associated with international clones of CRKp such as ST11, ST14, ST15, ST63, ST147, ST231, etc. his has aided in their rapid dissemination across boundaries [23].

Discordant phenotypic and genotypic resistance profile was observed in this study for aminoglycosides and folate pathway inhibitors, with higher genotypic as compared to phenotypic resistance. The discrepancy between phenotypic and genotypic resistance is a major shortcoming for the accuracy of genotypic antimicrobial resistance (AMR) prediction. Furthermore, differences in the expression of AMR genes may cause unjustified restriction of therapeutic options resulting in the use of less potent and more toxic substances [24]. The colistin susceptibility test results of 2 isolates were not in agreement with WGS results. Since ID/AST automated systems are known to produce variable in vitro colistin susceptibility test results, the joint Clinical and Laboratory Standards Institute (CLSI) and European Committee on Antimicrobial Susceptibility Testing (EUCAST) Polymyxin Breakpoints Working Group recommended broth micro dilution (BMD) method as the reference test for determining colistin susceptibility, in March 2016 [25]. Also, the phenotypic susceptibility test results of tigecycline cannot be considered reliable because as per EUCAST guidelines 2021, for Enterobacterales other than *Escherichia coli* and *Citrobacter koseri*, the activity of tigecycline varies from insufficient to variable [20]. Both CLSI and EUCAST have not defined any clinical breakpoints for tigecycline in case of *K. pneumoniae* [11,26].

Resistance to antimicrobials in *K. pneumoniae* is primarily acquired by horizontal gene transfer mediated by plasmids and/or mobile genetic elements [27]. The highly challenging MDR-Kp or CR-Kp isolates are mainly associated with the NDM or/and OXA-48-like carrying ST11, ST14, ST15, ST101, ST147, ST395 and ST231 in Asia and KPC harbouring ST258 in North America and Europe, respectively [28]. In the present study, all CRKp isolates showed phenotypic resistance to imipenem, meropenem and ertapenem, explained by the presence of *bla*_*OXA-48*_ like genes, *bla*_*NDM*_ genes, and a combination of ESBLs & inactivation of porins Ompk35/36 [29]. While 77.8% of these isolates carried *bla*_*OXA-48*_ like genes, 55.6% carried *bla*_*NDM-1/5*_, which supports the reported changing trends in carbapenem resistance [30].

Some of the lacunae of this study were small sample size and lack of a structured sampling framework. This study emphasizes the need for continued genomic surveillance of emerging CRKp and other MDR bacteria in Uttarakhand and neighbouring states of India. This in turn would help in generating critical information that can be used to assess the emergence, dissemination, and potential impact of important variants.

## Author statements

**Author statements:**
| Contributor role | Author 1 | Author 2 | Author 3 | Author 4 | Author 5 | Author 6 | Author 7 | Author 8 |
| --- | --- | --- | --- | --- | --- | --- | --- | --- |
| Conceptualization | Yes | Yes | Yes | Yes | Yes | Yes | - | Yes |
| Methodology | Yes | Yes | Yes | Yes | Yes | Yes | - | Yes |
| Software | - | Yes | - | - | - | - | - | - |
| Validation | Yes | Yes | Yes | Yes | Yes | Yes | - | Yes |
| Formal analysis | Yes | Yes | Yes | Yes | Yes | Yes | Yes | Yes |
| Investigation | Yes | Yes | Yes | Yes | Yes | Yes | - | Yes |
| Resources | Yes | Yes | Yes | Yes | Yes | Yes | Yes | Yes |
| Data curation | Yes | Yes | Yes | Yes | Yes | Yes | Yes | Yes |
| Original draft preparation | Yes | Yes | Yes | Yes | Yes | Yes | Yes | Yes |
| Visualisation | Yes | Yes | Yes | Yes | Yes | Yes | - | Yes |
| Supervision | Yes | Yes | Yes | Yes | Yes | Yes | - | Yes |
| Project administration | Yes | - | - | - | Yes | Yes | - | Yes |
| Funding | - | - | - | - | - | - | - | Yes (NIHR funded) |

## Ethical approval

Ethical approval was obtained vide letter #AIIMS/IEC/18/477 & KIMS IEC/S12-2017.

## References

1. Holt KE, Wertheim H, Zadoks RN, Baker S, Whitehouse CA, Dance D et al. Genomic analysis of diversity, population structure, virulence, and antimicrobial resistance in Klebsiella pneumoniae, an urgent threat to public health. Proc Natl Acad Sci U S A. 2015 Jul 7;112(27):E3574–81. doi: 10.1073/pnas.1501049112. Epub 2015 Jun 22. PMID: 26100894; PMCID: PMC4500264.

2. Dumitru IM, Dumitrascu M, Vlad ND, Cernat RC, Ilie-Serban C, Hangan A et al. Carbapenem-Resistant Klebsiella pneumoniae Associated with COVID-19. Antibiotics (Basel). 2021 May 11;10(5):561. doi: 10.3390/antibiotics10050561. PMID: 34065029; PMCID: PMC8151469.

3. Poirel L, Jayol A, Nordmann P. Polymyxins: antibacterial activity, susceptibility testing, and resistance mechanisms encoded by plasmids or chromosomes. Clin Microbiol Rev. 2017;30:557–96.

4. Karakonstantis S, Kritsotakis EI, Gikas A. Treatment options for K. pneumoniae, P. aeruginosa and A. baumannii co-resistant to carbapenems, aminoglycosides, polymyxins and tigecycline: an approach based on the mechanisms of resistance to carbapenems. Infection. 2020 Dec;48(6):835–851. doi: 10.1007/s15010-020-01520-6. Epub 2020 Sep 1. PMID: 32875545; PMCID: PMC7461763.

5. Chaudhry D, Tomar P. Antimicrobial resistance: The next big pandemic. Int J Community Med Public Health 2017;4:2632–6.

6. Gandra S, Joshi J, Trett A, Lamkang A, Laxminarayan R. Scoping Report on Antimicrobial Resistance in India. Washington, DC: Center for Disease Dynamics, Economics & Policy; 2017. Available from: http://www.dbtindia.nic.in/wpcontent/uploads/ScopingreportonAntimicrobialresistanceinIndia.pdf, accessed on June 20, 2021.

7. Perdigão J, Modesto A, Pereira AL, Neto O, Matos V, Godinho A et al. Whole-genome sequencing resolves a polyclonal outbreak by extended-spectrum beta-lactam and carbapenem-resistant Klebsiella pneumoniae in a Portuguese tertiary-care hospital. Microb Genom. 2019 Sep;7(6). doi: 10.1099/mgen.0.000349. PMID: 32234124.

8. World Health Organization. Global priority list of antibiotic-resistant bacteria to guide research, discovery, and development of new antibiotics; 2017. Available from: https://www.who.int/medicines/publications/WHO-PPL-Short_Summary_25Feb-ET_NM_WHO.pdf, [Last accessed on August 20, 2021].

9. Collee JG, Miles RS, Watt B. Tests for the identification of bacteria. In: Collee JG, Fraser AG, Marimon BP, Simmons A, editors. Mackie & McCartney Practical Medical Microbiology. 14th ed. New Delhi: Elsevier; 2007. p. 131–49.

10. Miles RS, Amyes SG. Laboratory control of antimicrobial therapy. In: Collee JG, Fraser AG, Marimon BP, Simmons A. editors. Mackie & McCartney Practical Medical Microbiology. 14th ed. New Delhi: Elsevier: 2007. p. 151–78.

11. Clinical and Laboratory Standards Institute. Performance Standards for Antimicrobial Susceptibility Testing; 31st Informational Supplement. CLSI Document M100. Wayne, PA: Clinical and Laboratory Standards Institute; 2021.

12. Prjibelski, A., Antipov, D., Meleshko, D., Lapidus, A., & Korobeynikov, A. (2020). Using SPAdes de novo assembler. Current Protocols in Bioinformatics, 70, e102. doi: 10.1002/cpbi.102.

13. https://www.protocols.io/view/ghru-genomic-surveillance-of-antimicrobial-resista-bpn6mmhe

14. Bowers JR, Lemmer D, Sahl JW, Pearson T, Driebe EM, Wojack B. KlebSeq, a diagnostic tool for surveillance, detection, and monitoring of Klebsiella pneumoniae. J Clin Microbiol 2016;54:2582–96.

15. Shankar C, Mathur P, Venkatesan M, Pragasam AK, Anandan S, Khurana S et al. Rapidly disseminating bla_OXA-232_ carrying Klebsiella pneumoniae belonging to ST231 in India: multiple and varied mobile genetic elements. BMC Microbiol. 2019 Jun 24;19(1):137. doi: 10.1186/s12866-019-1513-8. PMID: 31234800; PMCID: PMC6591861.

16. Navon-Venezia S, Kondratyeva K, Carattoli A. Klebsiella pneumoniae: a major worldwide source and shuttle for antibiotic resistance. FEMS Microbiol Rev. 2017;41(3):252–75.

17. Lee C-R, Lee JH, Park KS, Kim YB, Jeong BC, Lee SH. Global dissemination of carbapenemase-producing Klebsiella pneumoniae: epidemiology, genetic context, treatment options, and detection methods. Front Microbiol. 2016;7:895.

18. Momin MH, Liakopoulos A, Phee LM, Wareham DW. Emergence and nosocomial spread of carbapenem-resistant OXA-232-producing Klebsiella pneumoniae in Brunei Darussalam. J Global Antimicrob Resistance. 2017;9:96–99.

19. Mancini S, Poirel L, Tritten ML, Lienhard R, Bassi C, Nordmann P. Emergence of an MDR Klebsiella pneumoniae ST231 producing OXA-232 and RmtF in Switzerland. J. Antimicrob Mar 2018;73(3):821–23.

20. Wyres KL, Nguyen TNT, Lam MMC, Judd LM, van Vinh Chau N et al. Genomic surveillance for hypervirulence and multi-drug resistance in invasive Klebsiella pneumoniae from South and Southeast Asia. Genome Med. 2020 Jan 16;12(1):11. doi: 10.1186/s13073-019-0706-y. PMID: 31948471; PMCID: PMC6966826.

21. Shankar C, Venkatesan M, Rajan R, Mani D, Lal B, Prakash JAJ et al. Molecular characterization of colistin-resistant Klebsiella pneumoniae & its clonal relationship among Indian isolates. Indian J Med Res. 2019 Feb;149(2):199–207. doi: 10.4103/ijmr.IJMR_2087_17. PMID: 31219084; PMCID: PMC6563726.

22. Follador R, Heinz E, Wyres KL, Ellington MJ, Kowarik M, Holt KE et al. The diversity of Klebsiella pneumoniae surface polysaccharides. Microb Genomics. 2016;2:10.1099/mgen.0.000073.

23. Naha S, Sands K, Mukherjee S, Saha B, Dutta S, Basu S. OXA-181-Like Carbapenemases in Klebsiella pneumoniae ST14, ST15, ST23, ST48, and ST231 from Septicemic Neonates: Coexistence with NDM-5, Resistome, Transmissibility, and Genome Diversity. mSphere. 2021 Jan 13;6(1):e01156–20. doi: 10.1128/mSphere.01156-20. PMID: 33441403; PMCID: PMC7845606.

24. Kocer K, Klein S, Hildebrand D, Krall J, Heeg K, Nurjadi SBD. Pitfalls in genotypic antimicrobial susceptibility testing caused by low expression of bla_KPC_ in Escherichia coli. J. Antimicrob 2021; dkab267, https://doi.org/10.1093/jac/dkab267

25. European Society of Clinical Microbiology. EUCAST warnings Concerning Antimicrobial Susceptibility Testing Products or Procedures. EUCAST. Available from: http://www.eucast.org/ast_of_bacteria/warnings/. [Last accessed on 2021 Aug 09].

26. European Committee on Antimicrobial Susceptibility Testing (EUCAST). Breakpoint Tables for Interpretation of MICs and Zone Diameters. Version 11; 2021. Available from: http://www.eucast.org. [Last accessed on 2021 Aug 09].

27. Pitout JD, Nordmann P, Poirel L. Carbapenemase-Producing Klebsiella pneumoniae, a Key Pathogen Set for Global Nosocomial Dominance. Antimicrob Agents Chemother. 2015;59(10):5873–84.

28. Lai YC, Lu MC, Hsueh PR. Hypervirulence and carbapenem resistance: two distinct evolutionary directions that led high-risk Klebsiella pneumoniae clones to epidemic success. Expert Rev Mol Diagn. 2019 Sep;19(9):825–837. doi: 10.1080/14737159.2019.1649145. Epub 2019 Aug 1. PMID: 31343934.

29. Weng X, Shi Q, Wang S, Shi Y, Sun D, Yu Y. The Characterization of OXA-232 Carbapenemase-Producing ST437 Klebsiella pneumoniae in China. Can J Infect Dis Med Microbiol. 2020;2020:5626503. Published 2020 Jul 8. doi:10.1155/2020/5626503.

30. Ravikant, Kumar P, Ranotkar S, Zutshi S, Lahkar M. Spectrum β-lactamases (ESBL) in Escherichia coli isolated from a tertiary care hospital in North-East India. Indian J Exp Biol 2016; 54 : 108–14.

