## Supplementary material for "Molecular characterization of carbapenem resistant *Klebsiella pneumoniae* clinical isolates: Preliminary experience from a tertiary care teaching hospital in the Himalayas": Tables

| **ST** | **Test isolates**  **n (%)** | **KL-Loci**  **n (%)** | **O-Loci**  **n (%)** | **Predominant virulence factor genes**  **n (%)** | | | **Predominant plasmids***  **n(%)** | | | | | |
| --- | --- | --- | --- | --- | --- | --- | --- | --- | --- | --- | --- | --- |
|  |  |  |  | **Yersiniabactin** | **Aerobactin** | **rmpA2** | **IncFIB(pQil)** | **IncFIB(K)** | **Col44 0I** | **ColKP3** | **IncFII( K)** | **IncR** |
| 11 | 2(7.41) | KL105 - 2(100) | O2v2 - 2(100) | 2(100) | - | - | - | - | - | - | - | 2(100) |
| 14 | 8(29.63) | KL2 – 8(100) | O1v1 – 8(100) | 8(100) | - | - | - | 8(100) | - | 7(87.5) | 5(62.5) | 6(75) |
| 15 | 2(7.41) | KL35 – 1(50) ; KL2 – 1(50) | OL103 – 1(50); O1v1 – 1(50) | 1(50) | 1(50) | 1(50) | 1(50) | - | - | - | - | 1(50) |
| 43 | 2(7.41) | KL30 -2(100) | O1v1 – 2(100) | 2(100) | 2(100) | 2(100) | 2(100) | - | - | - | 2(100) | - |
| 147 | 3(11.11) | KL51 – 3(100) | O2v2 – 3(100) | 3(100) | - | - | - | - | - | - | - | - |
| 231 | 5(18.52) | KL51 – 5(100) | O1v2 – 5(100) | 5(100) | 4(80) | - | 5(100) | - | 5(100) | 5(100) | 4(80) | - |
| 395 | 2(7.41) | KL64 – 2(100) | O1v1 – 2(100) | 2(100) | - | - | 1(50) | - | - | 2(100) | - | - |
| 437 | 1(3.70) | KL36 – 1(100) | O4 – 1(100) | 1(100) | - | - | 1(100) | 1(100) | - | - | 1(100) | - |
| 3392 | 1(3.70) | KL64 – 1(100) | O1v1 – 1(100) | - | 1(100) | 1(100) | 1(100) | - | - | 1(100) | 1(100) | - |
| Novel | 1(3.70) | KL81 – 1(100) | OL101 – 1(100) | - | - | - | 1(100) | 1(100) | - | - | 1(100) | - |

Table-1: Summary of predominant virulence factor genes & plasmid repertoire across sequence types (STs) prevalent in CRKp test isolates

*Other plasmid replicons detected were Col(MG828), IncFIA, IncFII(pAMA1167-NDM-5), Col(pHAD28), IncFIB(pKPHS1), FIA(pBK30683), IncFIB(pNDM-Mar), IncHI1B(pNDM-MAR), IncC, IncX3, IncFII(pKPX1), IncN, IncFII, ColRNAI, IncFIB(AP001918), IncFIB(K)(pCAV1099-114), ColpVC, IncFIB(pQil)--IncFII(K), IncFIA(HI1) & repB_KLEB respectively.

Table-2: Phenotypic & genotypic antimicrobial resistance profile of CRKp test isolates

| **Antibiotic class** | **Phenotypic antimicrobial resistance**  **n (%)** | **Genotypic antimicrobial resistance**  **n (%)** |
| --- | --- | --- |
| **Aminoglycosides**  **(Amikacin, Gentamicin)** | 24 (88.9) | 26 (96.3) |
| **Fluoroquinolones**  **(Ciprofloxacin)** | 27 (100) | 27 (100) |
| **Penicillins**  **(Ampicillin)** | 27 (100) | 27 (100) |
| **Beta Lactam/Beta Lactamase Inhibitor combinations**  **(Amoxicillin/clavulanic acid, Piperacillin/tazobactam, Cefoperazone/sulbactam)** | 27 (100) | 27 (100) |
| **Cephalosporins**  **(Ceftriaxone, Cefuroxime axetil, Cefuroxime, Cefepime)** | 27 (100) | 27 (100) |
| **Folate pathway inhibitors**  **(Trimethoprim/sulfamethoxazole)** | 22 (81.5) | 25 (92.6) |
| **Polymyxins**  **(Colistin)** | 0 (0) | 1 (3.7) |
| **Glycylcyclines**  **(Tigecycline)** | 18 (66.7) | 0 (0) |

Table-3: Summary of predominant AMR genes across sequence types (STs) prevalent in CRKp test isolates

| **ST** | **Test isolates**  **n (%)** | **Predominant AMR genes*** | | | | | | | | |
| --- | --- | --- | --- | --- | --- | --- | --- | --- | --- | --- |
|  |  | **ESBL genes**  **n (%)** | | | | **Carbapenem genes**  **n (%)** | | | | **OmpK mutations**  **n (%)** |
|  |  | CTXM-15 | SHV-11 | OXA-1 | TEM-1 | NDM-1 | NDM-5 | OXA-181 | OXA-232 |  |
| 11 | 2(7.41) | 2(100) | 2(100) | - | - | - | 2(100) | - | - | - |
| 14 | 8(29.63) | 8(100) | 1(12.5) | 5(62.5) | 4(50) | 2(25) | 5(62.5) | 6(75) | 1(12.5) | 8(100) |
| 15 | 2(7.41) | 2(100) | - | - | 1(50) | 1(50) | 1(50) | - | - | 1(50) |
| 43 | 2(7.41) | 2(100) | 2(100) | - | 2(100) | - | - | 2(100) | - | 2(100) |
| 147 | 3(11.11) | 3(100) | 3(100) | - | 2(66.7) | - | 3(100) | 3(100) | - | 3(100) |
| 231 | 5(18.52) | 2(40) | - | - | - | - | - | - | 5(100) | 5(100) |
| 395 | 2(7.41) | 1(50) | 2(100) | 1(50) | 1(50) | - | 1(50) | - | 2(100) | 2(100) |
| 437 | 1(3.70) | 1(100) | 1(100) | - | 1(100) | - | 1(100) | 1(100) | - | - |
| 3392 | 1(3.70) | 1(100) | - | - | - | 1(100) | - | - | 1(100) | 1(100) |
| Novel | 1(3.70) | 1(100) | - | - | - | - | 1(100) | 1(100) | - | 1(100) |
